# CRISPR/Cas13a powered electrochemical microfluidic biosensor for nucleic acid amplification-free miRNA diagnostics

**DOI:** 10.1101/738617

**Authors:** Richard Bruch, Julia Baaske, Claire Chatelle, Mailin Meirich, Sibylle Madlener, Wilfried Weber, Can Dincer, Gerald Urban

## Abstract

Non-coding small RNAs, such as microRNAs, are becoming the biomarkers of choice for multiple diseases in clinical diagnostics. A dysregulation of these microRNAs can be associated to many different diseases, such as cancer, dementia or cardiovascular conditions. The key for an effective treatment is an accurate initial diagnosis at an early stage, improving the patient’s survival chances. Here, we introduce a CRISPR/Cas13a powered microfluidic, integrated electrochemical biosensor for the on-site detection of microRNAs. Through this unique combination, the quantification of the potential tumor markers microRNA miR-19b and miR-20a has been realized without any nucleic acid amplification. With a readout time of 9 minutes and an overall process time of less than 4 hours, a limit of detection of 10 pM was achieved, using a measuring volume of less than 0.6 µl. Furthermore, we demonstrate the feasibility of our versatile sensor platform to detect miR-19b in serum samples of children, suffering from brain cancer. The validation of our results with a standard qRT-PCR method shows the ability of our system to be a low-cost and target amplification-free tool for nucleic acid based diagnostics.

---

MicroRNAs (miRNAs) are composed of 18-25 nucleotides in their mature form and play an essential role as regulators of gene expression in many biological processes by binding to specific messenger RNA targets and promoting their degradation or translational inhibition^1,2^. In recent years, the popularity of miRNAs in research has been constantly growing, as the presence or dysregulation of distinct miRNAs can indicate specific medical conditions. For instance, in cancer research the up or down regulation of miRNA expression levels has been linked to certain cancer types, including lung cancer or brain tumors. This offers new possibilities for the detection and monitoring of such diseases, which is why miRNAs will continue to come in the focus of clinical diagnostics^3–5^.

Traditional methods of studying miRNA levels include microarrays, RNA sequencing methods and, as the current gold standard, the quantitative polymerase chain reaction (qPCR)^6^. Each of these methods, despite being powerful, have their limitations. Bulky and expensive equipment, intensive sample preparation or long turnover times limit most of them to well-equipped laboratories, restricting such screenings to developed countries. Therefore, there is a pressing need to develop easy-to-use, portable and amplification-free methods for miRNA detection at the point of care (POC). This will allow a fast, versatile and low-cost quantification of miRNAs not only in the western world, but also in resource-limited regions, where people do not have access to well-equipped laboratories and where cost efficiency is even more important.

Today, there are a few reports tackling these challenges, for example by improving current detection methods, using electrochemical assays^7,8^ or developing pH-based POC tests^9^. However, these approaches still need error-prone amplification steps prior to the miRNA detection, have rather tedious preparation steps or are limited by the design of suitable primers^10,11^. To address these issues, we combine microfluidics with an electrochemical signal readout to develop a sensitive (i.e. target amplification-free) and selective diagnostic test, while enabling a miniaturization for POC testing^12,13^. We further apply the newly discovered CRISPR/Cas13a technology, which is able of targeting almost any RNA, to our developed electrochemical biosensor, creating a powerful tool for miRNA diagnostics (Fig. 1a).

**Fig. 1.**
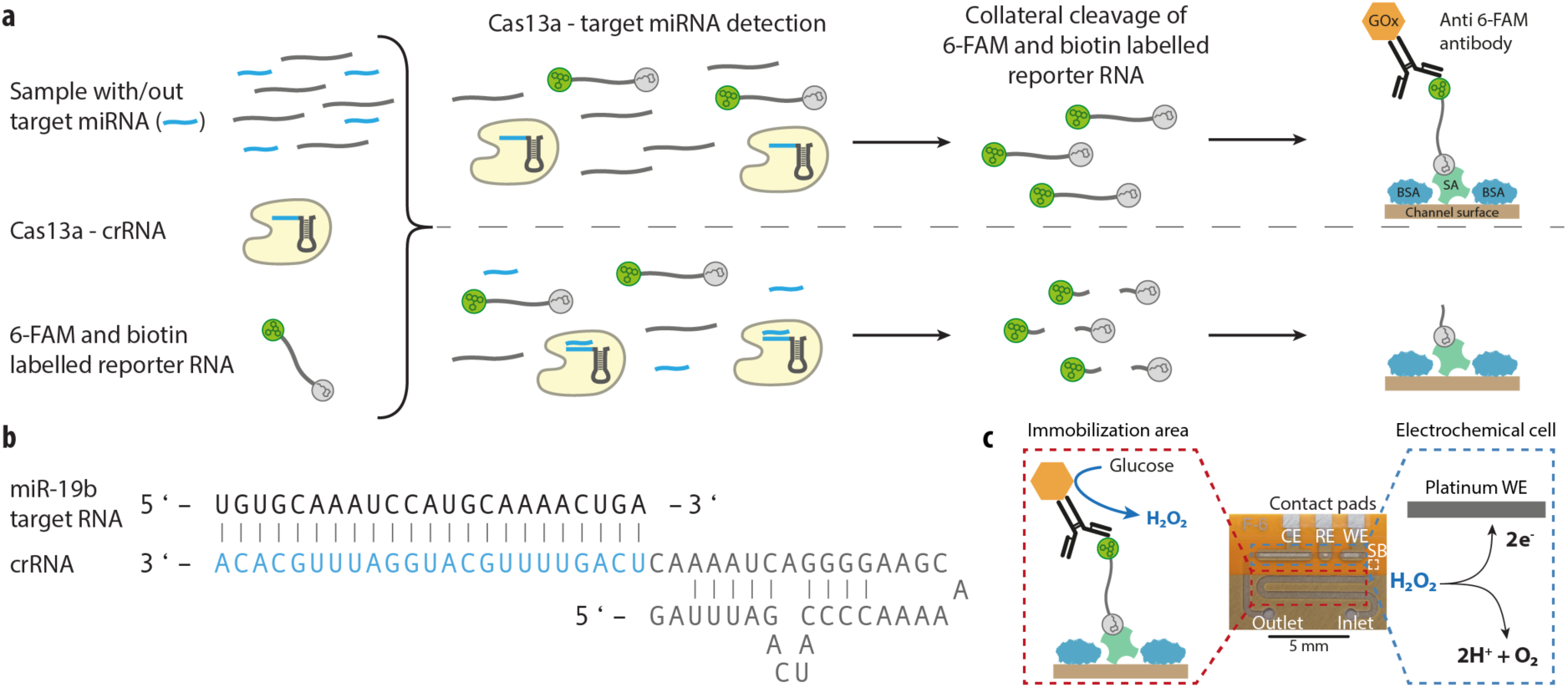
Combination of the CRISPR technology along with an electrochemical microfluidic biosensor for miRNA diagnostics. **a**, Schematic of the off-chip miRNA targeting, including the enzyme Cas13a, the target RNA, the target-specific crRNA and the biotin and 6-FAM labelled reporter RNA, which is immobilized after the cleavage process onto the streptavidin (SA) and BSA blocked channel surface. **b**, Schematic of the single stranded target miRNA, miR-19b, and the crRNA, where the complementary sequence is highlighted in blue. **c**, Working principle and photo of the microfluidic biosensor with its main elements, including the contact pads for the working, reference and counter electrode (WE, RE, CE) in the electrochemical cell (marked in blue) and the immobilization area for the assay preparation (highlighted in red), separated by the hydrophobic stopping barrier SB.

In the recent past, studies, employing clustered regularly interspaced short palindromic repeats (CRISPR) associated methods for the detection of nucleic acids, have been utilized using different Cas effectors^14^. Varying Cas effectors were used for targeting different nucleic acids, like the Cas9 effector for the detection of dsDNA^15–17^, along with the Cas12 for the detection of ssDNA^18–21^. The microbial CRISPR effector Cas13a (previously named C2c2), displays, in contrast to the other Cas effectors, a triggered cleavage capability of non-target ssRNAs in the surrounding^22,23^. Taking advantage of this characteristic, we replace synthetic nucleic acid amplification steps by a Cas13a driven signal amplification. The specificity of the detection is given by the need of a target-specific CRISPR RNA (crRNA) that guides the Cas13a to the RNA sequence of interest^23,24^ (Fig. **1**b). Upon recognition of the complementary RNA sequence, the cleavage ability of the enzyme is activated. The addition and subsequent cleavage of a reporter RNA (reRNA) can thus be used for a quantitative readout of RNA levels^25^. Since after the activation of the Cas/crRNA complex the collateral cleavage mechanism is, in contrast to other Cas effectors, an ongoing process, the amount of cleaved reRNA is a time depending progress, resulting in a self-amplification. These two specific characteristics of the Cas enzyme enable a highly selective and sensitive detection. So far, Cas13a was used for the optical detection of virus RNAs or even for the detection of plant genes^26^, whereas no research work exists up to now, combining CRISPR/Cas13a with an electrochemical detection method for miRNA diagnostics^23–25,27–31^. Here, we present the CRISPR technology on a microfluidic biosensor, electrochemically measuring miRNA levels of the potential brain tumor marker miR-19b in serum samples from patients, suffering from brain cancer.

For the detection of miRNAs, a low-cost and easy-to-use electrochemical biosensor was manufactured, using the dry film photoresist (DFR) technology^32,33^. By stacking multiple developed DFR foils onto a platinum patterned polyimide substrate, the microchannel and the electrodes are realized. The microfluidic channel thereby consists out of two distinct sections: (i) an immobilization area and (ii) an electrochemical cell, separated by a hydrophobic stopping barrier (Fig. 1c). The immobilization area is filled and functionalized through an inlet by capillary forces, until the flow reaches the stopping barrier. This allows an easy handling of the chip, combined with an automatic metering (< 0.6 µl) of the introduced biomolecules and, in addition, prohibits a contamination of the electrodes. In the electrochemical cell, the detection of enzymatically produced hydrogen peroxide (H_2_O_2_) takes place, using a three-electrode setup, comprising a platinum working and counter electrode together with a silver/silver chloride reference electrode.

For the CRISPR/Cas13a powered miRNA detection, the surface of the immobilization area of the biosensor is pre-functionalized by applying streptavidin to the chip inlet, which is followed by a blocking step, using bovine serum albumin (BSA). After each incubation step, a washing step is performed, where the microchannel is flushed with wash buffer. By applying a vacuum to the channel inlet, all unbound biomolecules are removed, without contaminating the electrochemical measurement cell. For the activation of the Cas13a and the subsequent cleavage process, the enzyme Cas13a is mixed in a standard microcentrifuge tube with its target-specific crRNA, a biotin and 6-FAM labelled reRNA and the sample of interest, potentially containing the target miRNA. The mixture is incubated in the standard tube for 1 to 24 hours at 37 °C, where the Cas13a forms a complex with the target-specific crRNA. In the presence of target miRNAs, the Cas13a gets activated, resulting in a collateral cleavage of the surrounding reRNA.

After the off-chip targeting of the miRNAs, the mixture is subsequently applied to the pre-functionalized microfluidic chip, where the cleaved and non-cleaved reRNAs, depending on the quantity of active enzymes, bind to the immobilized streptavidin. Following a washing step, anti-fluorescein antibodies coupled to glucose oxidase (GOx), which are only capable of binding to the uncleaved reRNAs, are introduced, enabling an enzymatic reading of the assay (Fig. 1a).

For the assay readout, the biosensor chip is placed into a custom-made holder, allowing the electrical connection to the potentiostat and the fluidic connection to a syringe pump. By pumping a glucose solution through the microfluidic biosensor, GOx catalyzes its substrate, producing H_2_O_2_, which is amperometrically detected in the electrochemical cell. For a further signal amplification, a fully automated so-called stop-flow protocol is used^34^. Herein, the pump is programmed so that the flow of the glucose solution is stopped for a certain period of time, where H_2_O_2_ is produced by the enzyme and accumulates inside the immobilization area. The pump automatically restarts the flow, the enriched H_2_O_2_ concentration is flushed over the working electrode, which results in a current peak with a specific peak charge. The gained amperometric signal is directly proportional to the amount of immobilized GOx, bound to the uncleaved reRNA and, therefore, inversely proportional to the concentration of target miRNA in the sample.

The assay’s sensitivity was improved through optimizing each assay component in terms of incubation time and concentration (Supplementary Fig. 2-10). Furthermore, to overcome the diffusion limitation of the reagents, a dynamic cleavage process, through shaking the mixture solution, while incubating at 37 °C, was implemented. As it became apparent, the shaking of the solution hindered the catalytic activity of the Cas13a (Supplementary Fig. 11), why further on a static cleavage process was chosen. As the crRNA needs to bind to the Cas effector in order to initiate the cleavage activity, the shaking might hamper this binding process or even promotes the loss of already bound crRNA.

By preparing solutions, containing different concentrations (100 fM to 10 nM) of the miRNA miR-19b and miR-20a, incubating them at 37 °C and applying them to the functionalized channel, different calibration curves were recorded (Fig. 2, Supplementary Fig. 15 and 16). By fitting the measured data points to a 4 or 5-parametric sigmoidal curve, a limit of detection (LOD) of 10 pM, equivalent to an amount of 500 amole of miRNA, was achieved for the miR-19b within 3 hours incubation time and an overall inter-assay coefficient of variation of less than 10% (Fig. 2b). As the cleavage of the reporter RNA is a time dependent process, the prolongation of the incubation increases the amount of cleaved RNA and, therefore, reduces the amperometric signal (Supplementary Fig. 14). To decrease the limit of detection furthermore, a calibration curve with an extended incubation time of the mixture solution of 7 hours (Supplementary Fig. 15) and 24 hours is recorded (Fig. 2c). Contra intuitive, we could not observe an improvement of the LOD for a 7-hour cleavage time. In our opinion, this is due to the very low concentration of the target miRNA and the therewith resulting lack of active Cas13a enzymes in the solution. This leads to an unsatisfactory total catalytic activity of the effector with a very low reaction rate, which is not sufficient for a further significant signal reduction at very low concentrations of the target miRNA within 7 hours of incubation. Comparing the different cleavage times in terms of sensor performance, a minimum LOD of 2 pM was achieved after a 24 hour incubation, while a maximum dynamic range of roughly 2 order of magnitudes can be seen, at a cleavage time of only 1 hour, along with a LOD of 18 pM (Fig. 2a). In general, the longer the cleavage time, the lower the limit of detection and the lower the dynamic range of the resulting calibration curve (Supplementary Table 6). With that, our CRISPR/Cas13a assay powered biosensor is able to reach the stated LOD of 10 pM after 3 hours, showing its capability to detect picomolar concentrations of miRNAs, without any pre-amplification procedure of the target miRNA, while consuming reagent volumes less than 0.6 µl per incubation.

**Fig. 2.**
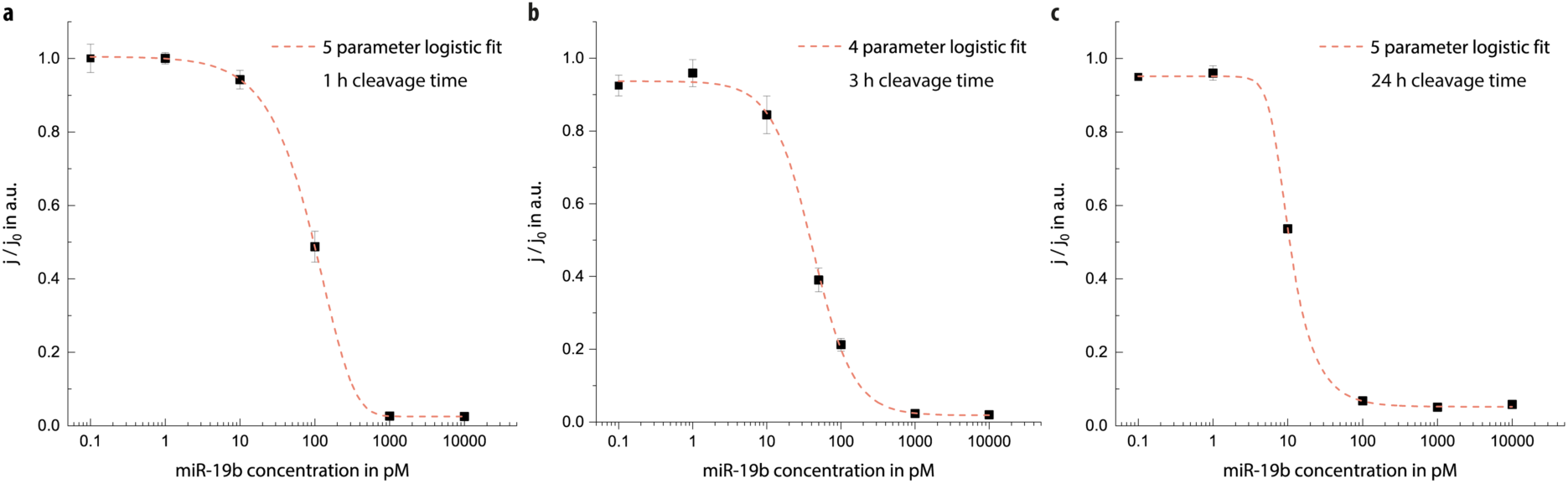
Time dependency of the CRISPR based off-chip miRNA targeting. **a**, Calibration curve of the off-chip cleavage scenario for 1 hour, using the miR-19b as a target miRNA. The reporter RNA (reRNA) is applied at a concentration of 250 nM, for the enzyme Cas13a and the crRNA a concentration ratio of 1:2 and 1:4 with respect to the reRNA is used, respectively. The results are fitted with a 5-parametric logistic fit, gaining a limit of detection of 18 pM with an inter-assay coefficient of variation (CV) of below 9%. With n = 4 replicates, error bars represent ± SD. **b**, Off-chip calibration curve, as described in Fig. 2a, using the miR-19b as target miRNA. The incubation time for the cleavage process was 3 hours. The results are fitted with a 4-parametric logistic fit, resulting in a limit of detection of 10 pM, along with a CV of below 10%. All error bars represent ± SD of n = 8 replicates. **c**, Calibration curve of the off-chip cleavage method as in Fig. 2a, using the miR-19b as target miRNA. The incubation time for the cleavage process was 24 hours. The results are fitted with a 5-parametric logistic fit, resulting in a limit of detection of 2 pM and a CV of below 8% in the dynamic range. All error bars represent ± SD of n = 4 replicates.

In order to validate the feasibility of our sensor concept for the detection of low miRNA concentrations in clinically relevant specimens, four serum samples of children, suffering from medulloblastoma were tested. Medulloblastoma is an aggressive embryonal tumor of the cerebellum/forth ventricle, which is characterized by an inhibition of apoptosis and an increased proliferation, characteristics caused by altered expression patterns in miRNAs^35^. This kind of brain tumor is one of the most common pediatric malignant tumors, often showing an aggressive progression^36^. It has been reported that, compared to healthy children, the concentration of the miR-17 ∼ 92 cluster family, containing the here employed biomarkers miR-19b and miR-20a, are upregulated for this tumor type^36–38^. Therefore, samples of patients, showing different clinical conditions, were chosen. Patient 1 and 3 achieved a complete remission of the tumor, whereas patient 2 and 4 developed a progressive state within a few months, which suggests that the patients 2 and 4 should have a significantly elevated miR-19b level. For the biochip measurements, the total RNA from the serum samples was isolated, using a commonly available RNA purification kit. The purified RNA was mixed with the Cas13a, its crRNA, specific to the miR-19b, and incubated at 37 °C. Consequently, the on-chip incubation and signal readout were performed as illustrated in Fig. 1c.

To confirm our biochip measurements, a standard quantitative real-time polymerase chain reaction (qRT-PCR) method was executed to measure the miRNA levels of the miR-19b as well. The measurements were performed from the same samples with a similar RNA purification technique. A serum medley of a pool of 20 healthy patients was used as a control measurement and treated the same way as the samples of the four patients, suffering from medulloblastoma. The results of both measurement methods show a good agreement within all four patient samples, especially by taking the nonlinearity of the measurement setup and the fact that the qRT-PCR cannot distinguish between miR-19a and 19b into consideration (Fig. 3a). Besides being a powerful method for the detection of nucleic acids, qRT-PCR methods normally require sophisticated systems and sample preparation, like for the reverse transcription of the cDNA, a long hands-on time and well-trained operators. In comparison with that, our method for miRNA detection simplifies the procedure and enables thereby the detection in a low-cost and easy manner, without the use of complex signal amplification methods or special trained personnel.

**Fig. 3.**
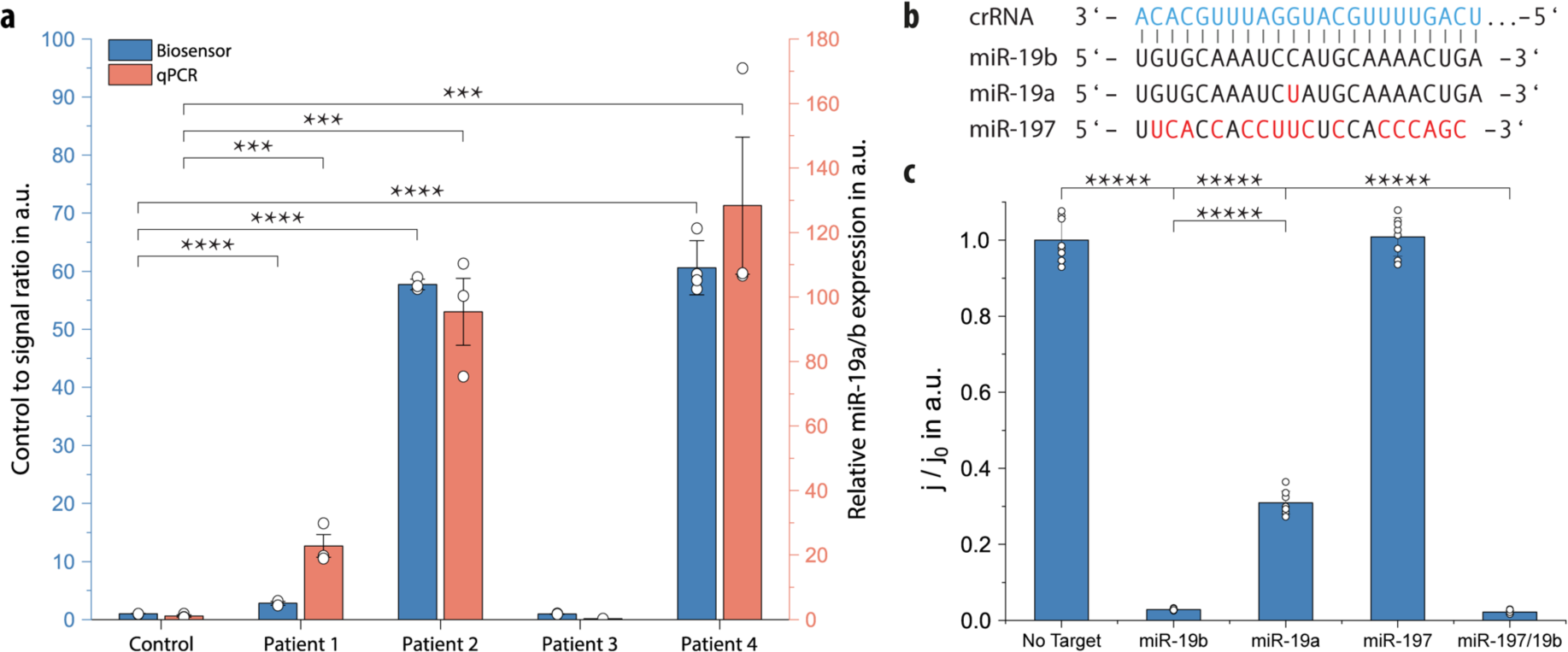
CRISPR based off-chip miRNA targeting. **a**, Validation of our measurement principle, using four serum samples from children, suffering from medulloblastoma and a control sample of a pool of 20 healthy patients. For the biosensor measurements, error bars represent ± SD of n = 4 replicates; error bars of the qPCR measurement represent ± SEM of n = 3 measurements; two-tailed Student’s t-test; ***P < 0.001; ****P < 0.0001. **b**, Schematic of the crRNA target region and the target sequences used for the detection. The mismatches in the target sequences are highlighted in red. **c**, Comparison of the cleavage efficiency of different miRNA targets, completely matching the complementary crRNA sequence (miR-19b), having a single mismatch (miR-19a) or do not match to a greater extend (miR-197). Applied concentration of 1 nM for each miRNA with n = 8 replicates, two-tailed Student’s t-test; *****P < 0.00001; error bars represent ± SD.

For an accurate diagnosis, a highly selective detection of the target miRNA is crucial. To investigate the biosensor’s specificity, target miRNAs, differing from the miR-19b by a single nucleotide (miR-19a) or several nucleotides (miR-197), were tested (Fig. 3b). For this, concentrations of 1 nM of the miR-19b, miR-19a, miR-197 as well as a mixture of the miR-197 and miR-19b were added to the sample solution, containing the Cas13a and the crRNA, complementary to the miR-19b. For the miR-197, the Cas13a does not show any cleavage ability, as the hybridization of the target miRNA to the crRNA is not sufficient to activate the enzyme’s catalytic activity. Comparing the miRNA miR-19b with the miR-19a, containing a single-base variation in the middle of the sequence, the biosensor enables a single-base mismatch detection within a significance greater of P < 0.00001 at low concentrations (Fig. 3c). As both miRNAs (miR-19a and 19b), together with the miRNAs miR-17, miR-18a, miR-20a and miR-92a, are part of the miR-17 ∼ 92 cluster family^37^, upregulated in patients, suffering from medulloblastoma, the exact distinction of these miRNAs is not compulsory necessary for an accurate diagnosis of this type of tumor. Additionally, to enhance the specificity of a future system, synthetic mismatches within the crRNA can further increase the signal, while the Cas13a is able to distinguish between single-base mismatches, as reported by Gootenberg and colleagues^27^.

For a more versatile and easy-to-handle biosensor, we also envision a “one-for-all” sensor chip, containing a complete pre-immobilized assay. In contrast to the off-chip miRNA targeting, where the activation of the Cas13a and the cleavage process were performed off-chip in a standard tube, the “one-for-all” sensor chip allows the activation and the cleavage to be done on-chip, while incubating. For the assay, an anti-biotin antibody instead of streptavidin is used as a surface coating for the binding of the biotin labelled reRNA. By using streptavidin to couple the reRNA to the surface for the on-chip cleavage, we believe that the Cas13a was sterically hindered, resulting in an inefficient cleavage process of the immobilized reRNA (Supplementary Fig. 18). To overcome this issue, a polyclonal antibiotin antibody is used, creating a greater distance to the channel surface and thus, providing a better accessibility of the Cas13a to the reRNA (Fig. 4a).

**Fig. 4.**
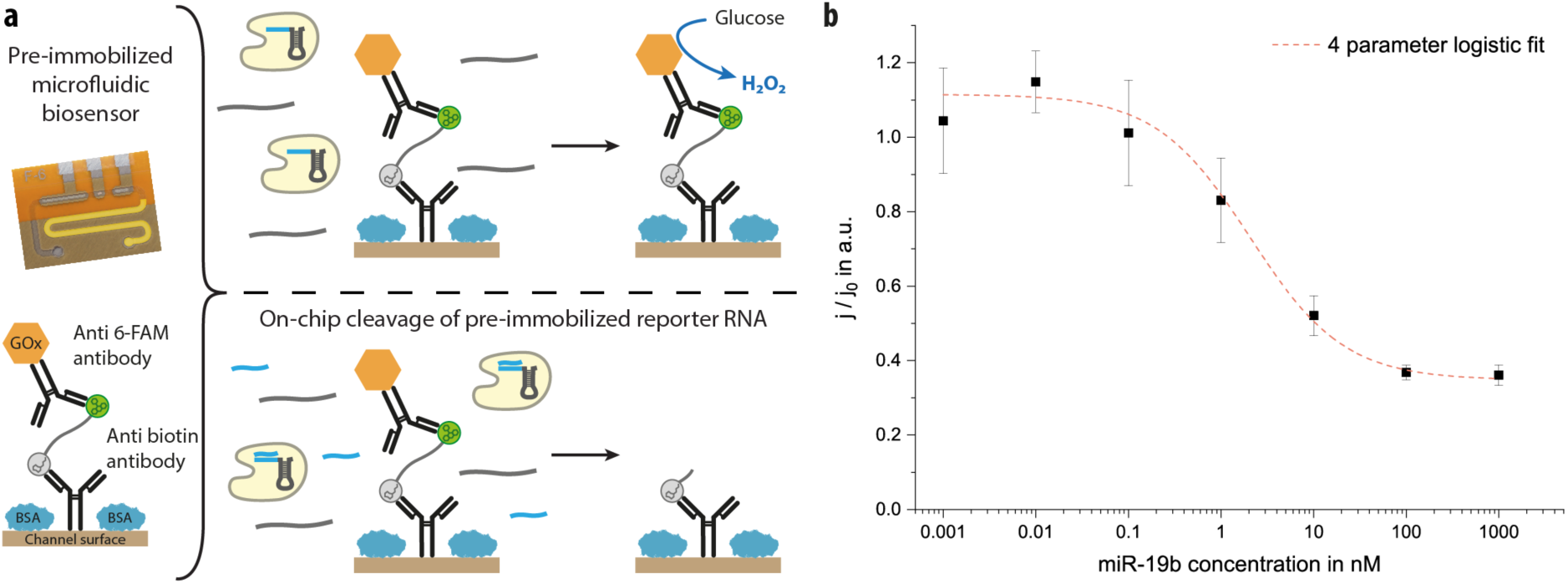
The on-chip miRNA detection: a “one-for-all” biosensor. **a**, Schematic, illustrating the on-chip cleavage procedure for samples with or without target miRNAs (blue). On the left, the sensor is depicted, showing the immobilization area, highlighted in yellow with the pre-immobilized assay, comprising an anti-biotin antibody surface coating along with a BSA blocking. The biotin and 6-FAM labelled reRNA is immobilized to the anti-biotin antibody, where the GOx labelled anti-fluorescein antibody binds to the reRNA. On the right, a mixture, containing the crRNA/Cas13a complex and a sample with and without target miRNAs (blue), is introduced into the biosensor to the completely pre-immobilized assay. The target activated enzyme cleaves the bound reRNA and thus, enables the removal of the GOx labelled antibody, resulting in a reduced amperometric signal. **b**, Calibration curve of the “one-for-all” biosensor for an on-chip miRNA detection, using the miR-19b as a target miRNA. The results are fitted with a 4-parametric logistic fit, gaining a limit of detection of 2.2 nM with an interassay coefficient of variation of below 15%. With n = 4 replicates, error bars represent ± SD.

All assay components, including anti-biotin antibodies for the surface coating, the biotin and 6-FAM labelled reRNA and the GOx labelled anti-fluorescein antibody, are immobilized in the biosensor’s immobilization area. For the detection of any miRNA, only the mixture of the sample, potentially containing the target miRNA and the target-specific crRNA/Cas13a complex need to be introduced to the microchannel and incubated for at 37 °C. While the mixture is incubated, the enzyme cleaves, in presence of any target miRNA, the immobilized reRNA and thus, remobilizes the GOx labelled antibodies. Following a washing step to remove all unbound GOx labelled antibodies, the assay readout can be performed by introducing the glucose solution (Fig. 4a). The produced H_2_O_2_ results, as before, in a current density proportional to the amount of bound GOx and is reverse proportional to the concentration of target miRNA in the sample solution.

For a first examination of the on-chip detection with the “one-for-all” biosensor, a calibration curve was performed. Solutions, containing different concentrations (1 pM to 1 µM) of the miRNA miR-19b, were prepared and introduced to the completely functionalized microchannel and incubated at 37 °C for 3 hours. Furthermore, to increase the cleavage capacity of the Cas13a, the solution is exchanged hourly during the incubation process. After the amperometric readout, the data points were fitted to a 4-parametric logistic curve, resulting in a LOD of 2.2 nM with an overall inter-assay coefficient of variation of less than 15% (Fig. 4b). This proves the feasibility to transfer the complete assay to the biosensor for an on-chip detection of low amounts (< 5 fmole) of any miRNA at the point of need in a low-cost manner, without the need of expensive equipment. Nevertheless, as the gap in the LOD values of the off-chip miRNA targeting *versus* the on-chip detection is evident, the on-chip detection needs further investigation and optimization.

## Discussion

Since the discovery of the CRISPR class 2 effector Cas13a in 2015^14^, different approaches for the utilization of the effector for biosensing applications were presented in literature^27–29,39^. However, no work has been reported so far for a CRISPR powered assay, along with an electrochemical detection method. Here, we have utilized the unique combination of a microfluidic, electrochemical biosensor with the powerful CRISPR/Cas13a technology for the quantification of low concentrations of miRNAs in serum specimens. Electrochemical, microfluidic biosensors have shown in the past their capability of detecting low concentrations of analytes in clinically relevant samples on an easy-to-fabricate integrated sensing platform, without the need of extensive instrumentations^40^. In particular, we use the CRISPR/Cas13a technology for a self-amplification upon recognition of the target miRNA, combined with a fully automated microfluidic stop-flow protocol for a further amplification of the electrochemical signal. By this extraordinary combination, our detection method requires no nucleic acid amplification prior to the detection procedure and, therefore, no additional costly reagents or specialist equipment, including trained personnel is needed.

Moreover, amplification-free CRISPR powered biosensing methods are rare and up till now only one Cas9 based approach exist for an amplification-free detection of dsDNA on a so-called CRISPR-Chip, based on a graphene field-effect transistor (gFET)^16^. For the detection of dsDNAs, several incubation steps lead to the final functionalized CRISPR-Chip, having a readout time of down to 15 min, comparable to our stop-flow amplified readout process of roughly 9 min (Supplementary Fig. 17). The CRISPR-Chip aims for the detection of gene mutations for the diagnostic of Duchenne muscular dystrophy associated mutations in the human genome^41,42^. For this, they are able to use the whole human genome (1.9 x 10^12^ g mol^-1^), which leads to an amplification-free detection limit of 1.7 fM (3.3 ng µl^-1^). By taking the molecular weight of our employed miRNA miR-19b (7.3 x 10^3^ g mol^-1^) for a similar calculation, a LOD of our biosensor of 73 fg µl^-1^ is achieved. For transferring the CRISPR-Chip to be able to detect RNAs, the effector Cas13a could be used as well. As the signal generation of the gFET is based on the amount of adsorbed and interacting charged molecules, the LOD of the CRISPR-Chip of 3.3 ng µl^-1^ would most likely stay the same, leading to a LOD of roughly 0.45 µM for the here employed miRNA miR-19b.

One major drawback of many reported diagnostic tools for miRNA detection is the extensive preparation of patient samples for the quantification of the target analyte. Often isolation and purification kits are needed, to extract the desired target without any other interfering biomolecules. Even though, our electrochemical biosensor needs one kit as well for the RNA isolation from serum, it is a commonly available one with a wide range of application fields. In general, the abandonment of such purification kits would drastically improve CRISPR based detection methods, allowing an easier handling and a faster gauging of the desired analyte.

Despite the powerful detection of other CRISPR based detection methods, the employed chips are often highly complex and requires multiple fabrication steps. The employed gFET for the CRISPR-Chip, for example, is based on silicon fabrication techniques, which normally requires sensible clean room stages, making it a powerful, bust costly detection tool. Our employed biosensor is based on polyimide and dry film photoresist layers, which are fairly easy to fabricate. On the other side, one clean room step is also needed for realizing the platinum electrodes of the electrochemical cell of our biosensor, which accounts of roughly two-thirds of the total fabrication costs. The employed reagents, like antibodies, are partly costly as well, resulting in an all-over price of roughly 0.75 € per functionalized biosensor, manufactured under research conditions (Supplementary Table 4). In general, our sensor device or silicon-based systems can be produced in standard industrial facilities, which will reduce the manufacturing price per chip drastically.

In this work, we have shown that our CRISPR/Cas13a powered biosensor efficiently and accurately detects miRNAs in small sample volumes, without the need of any nucleic acid amplification step. The clinical potential of our system was successfully evaluated by measuring different serum samples of pediatric patients, suffering from medulloblastoma in different stages. By examining the same samples with a standard qRT-PCR method, the findings of our biosensing chip were validated within a good agreement. These results indicate that our biosensor chip is capable of diagnosing miRNA related diseases, like brain tumors, by bypassing of any pre-amplification step. We furthermore overcome limitations, including the restrictions of common primer designs, extending this technique to short target miRNAs previously difficult to detect and with the obtained minimal detection limit of 2 pM, we are able to measure miRNAs in the clinical range of circulating miRNAs in serum^43^.

Although we are rivaling other Cas13a powered optical detection methods for longer RNAs (i.e. SHERLOCK) by a factor of five^27^ prior to their nucleic acid amplification step, as reported by Li and colleagues^24^, other electrochemical methods for the detection of miRNAs exist, having lower LODs as our presented approach^44–54^. The basic working principle of most of these electrochemical sensors is the immobilization of a (labelled) nucleic acid sequence, complementary to the target sequence, onto a screen-printed (gold) electrode, which enables an amplification of the signal, resulting in low LODs. Besides the higher material values used in these sensors, this also indicates a much higher hands-on time for the incubation process for the functionalization of the electrode, as each electrochemical chip must be target-specifically pre-treated before the desired miRNA can be detected. In contrast to that, our biosensor chip can be easily pre-functionalized and stored for up to nine months under refrigerated conditions (Supplementary Fig. 13) without the need of any target specific pre-treatment prior to the detection of the desired miRNA, compared to other electrochemical systems, having a stability in the range of days or weeks^55^.

By presenting the possibility of transferring the complete assay for the miRNA detection on our “one-for-all” biosensor chip, it shows the feasibility to detect any desired miRNA, requiring only the input of the Cas13a/crRNA – target mixture. Although the “one-for-all” biosensor needs further improvements in terms of sensitivity and reproducibility, it does show the practicability of this system for an easy-to-use setup, reducing many incubation steps, while decreasing the sample-to-result time. Overall, the unique fusion of the CRISPR/Cas13a technology with our microfluidic biosensor, combined with an electrochemical readout, allows the detection of the desired miRNA with high sensitivity and selectivity with a short hands-on time. This pioneers the way for a new generation of on-site miRNA diagnostic tools.

Future work will be the further reduction of the LOD by implementing different strategies, like an immobilization of the Cas/crRNA complex. With that, the target sample can be flushed over the immobilized complex, while the target sequence will bind to the complex and activates the catalytic cleavage mechanism of the Cas effector. Employing this strategy to the developed biosensor, a strong reduction in the LOD should be noticeable. Besides an optimization of the immobilization strategy, an additional electrochemical signal amplification via a so-called redox cycling by interdigitated electrode arrays (IDAs) can be combined with our microfluidic stop-flow measurement protocol. In our previous published works, signal amplifications of more than 150-times is achieved, employing nanogap IDAs with 100 nm gap size^56–58^. With a combination of this technique and our CRISPR based biosensor, the LOD of our system could be reduced to less than 100 fM.

Furthermore, a future multiplexed version of our sensor system, comprising several incubation channels, could allow the detection of multiple miRNAs from one single sample, resulting in a more accurate diagnosis of a certain disease^59^. This will not only allow us to detect one specific miRNA, but also enables the realization of a positive (by the use of a specific target concentration) and negative (employing no target) built-in control for each measurement. Such a CRISPR/Cas powered multiplexed biosensor, being able of measuring up to 8 miRNAs (Supplementary Fig. 20), will also deliver a more complete picture of the patient’s disease, for which nowadays expensive and lab-intensive equipment is needed.

## Methods

Any supplementary information is available in the Supplementary.

## Acknowledgments

The authors would like to thank the German Research Foundation (DFG) for the funding of this work under grant numbers UR 70/10-01, UR 70/12-01, WE 4733/4 and EXC-294 (BIOSS). We also thank Dr. Maximilian Hörner for providing technical advice for using ÄKTA chromatography systems, Dr. Nils Schneider for helping us with RNA concentration measurements and we would like to thank Dr. Chryssa Grylli for taking the patient blood samples.

## Author contributions

R.B., J.B. and C.D. designed the experiments. R.B. performed the biosensor experiments and analyzed the data. S.M designed and performed the qRT-PCR measurements. J.B. performed the preliminary enzyme work. R.B., J.B. and C.D. wrote the manuscript. C.C. and M.M gave technical advice. C.D. conceived and managed the project and W.W. and G.U. supervised the work.

## Ethics

The research was authorized by the Ethic Commission of the Medical University of Vienna (1244/2016).

