## Supplementary information for "CRISPR/Cas13a powered electrochemical microfluidic biosensor for nucleic acid amplification-free miRNA diagnostics"

### Materials and Methods:

**Plasmids used during this study:**

pC013[1] was purchased from addgene (#90097).

**LwCas13a protein purification:**

LwCas13a was purified similar to Gootenberg et al.[1] .Briefly, LwCas13a bacterial expression vector was transformed into Rosetta2 (DE3) pLysS competent cells. Starter culture was grown over night in Terrific Broth growth media (12 g $l^{-1}$ BD BactoTM tryptone (BD Biosciences, cat. no.: 211705), 24 g $l^{-1}$ BD BactoTM yeast extract (BD Biosciences, cat. no.:288610), 9.4 g $l^{-1}$ $K_2HPO$ (Roth, cat. no.: 6875), 2.2 g $l^{-1}$ $KH_2PO_4$ (Roth, cat. no.: 3904)) supplemented with 1:1,000 dilution of Ampicillin (stock: 100 mg $ml^{-1}$) and Chloramphenicol (stock: 36 mg $ml^{-1}$). Starter culture was used to inoculate 4 l of TB for growth at 37 °C and 150 rpm until OD600 of ~0.7. Protein expression was subsequently induced with 500 µM IPTG (final concentration) and cells were placed at 18 °C for at least 16 h for protein expression. Cells were centrifuged at 5,200 × g for 15 min at 4 °C. Cell pellet was frozen in liquid nitrogen and stored at −80 °C for further use.

All following steps were performed at 4 °C. Cell pellet was thawed on ice and resuspended in lysis buffer (20 mM Tris-HCl, 500 mM NaCl, 1 mM DTT, pH 8.0). Homogenize at 4 °C and subsequently cells were disrupted using a French press (APV 2000, APV Manufacturing) at a maximum of 1,000 bar. Lysate was spun down for 1 h at 30,000 × g at 4 °C. Cleared supernatant was applied to a total of 4 ml of StrepTactin Sepharose (IBA, cat. no.:2-4030-002), incubated for 1 h at 4 °C with rotation, followed by washing of the protein-bound resin with lysis buffer. After, the resin was washed, it was resuspended in SUMO digest buffer (30 mM Tris-HCl, 500 mM NaCl, 1 mM DTT, 0.15% Igepal (NP-40), pH 8.0). To cleave protein off resin, 250 units of SUMO protease (ThermoFisher Scientific, cat. no.: 12588018) were added and incubated over night at 4 °C with rotation. Protein eluate was isolated by filtering StrepTactin beads. For further cation exchange, protein was concentrated using 20 ml Spin-X UF Concentrator (10 kDa MWCO, omnilab, cat. no.: CORN431488) and buffer exchanged to elution buffer (containing 130 mM NaCl, 20 mM Tris-HCl, 1 mM DTT, 5% glycerol, pH 8.0), using Pierce Dextran desalting columns (ThermoFisher Scientific, cat. no.: 43233). Concentrated protein was loaded onto a 5 ml HiTrap SP HP cation exchange column (GE Healthcare Life Sciences, cat. no.: 17115201) via ÄKTAxpress (GE Healthcare Life Sciences) and eluted over salt gradient from 130 mM to 2 M NaCl in elution buffer. Fractions containing protein were tested for presence of LwCas13a by SDS-PAGE. Protein containing fractions were pooled, concentrated as before, and subsequently loaded on gel filtration column (Superdex® 200 10/300 GL, GE Healthcare Life Sciences, cat. no.: 17-5175-01) on an ÄKTAprime plus (GE Healthcare Life Sciences) buffered with S200 buffer (10 mM HEPES, 1 M NaCl, 5 mM $MgCl_2$, 2 mM DTT, pH 7.0). The resulting protein containing fractions were analyzed with SDS-PAGE for presence of LwCas13a. Those fractions were pooled and buffer exchanged to storage buffer (600 mM NaCl, 50 mM TrisHCl, pH 7.5, 5% glycerol, 2 mM DTT). Concentration was determined using Bradford assay (Bio-Rad, cat. no.: 500-0006) with bovine serum albumin (Sigma Aldrich, cat. no.: 05479) as standard. Protein was aliquoted at 4 °C and subsequently stored at −80 °C.

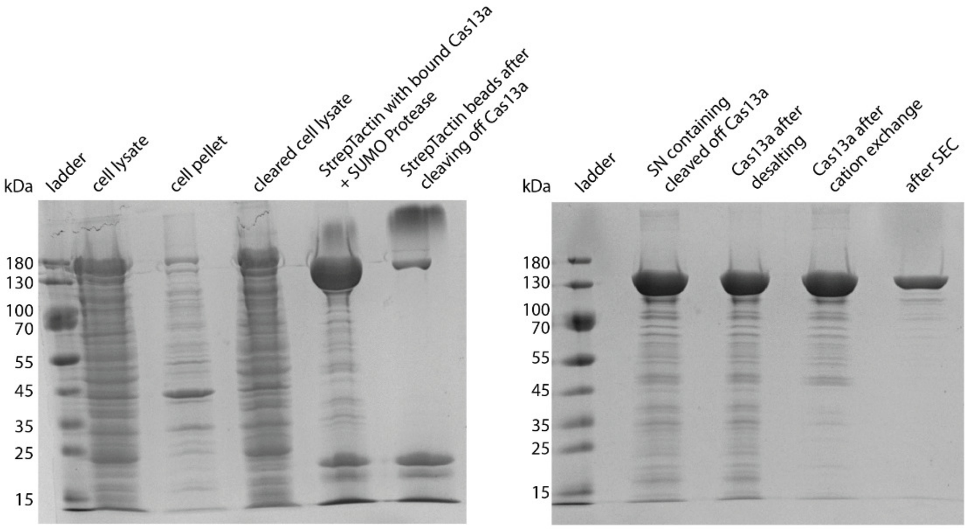

**Fig. 1 |** LwCas13a purification. Coomassie blue stained acrylamide gels of LwCas13a stepwise purification. Final purified LwCas13a after size exclusion chromatography (SEC).

**Nucleic acid targets and crRNA preparation:**

All target miRNAs (miR-19b, miR-19a, miR-20a and miR-197) and all biotin and 6-FAM labeled reporter RNA and DNA (reRNA_14b, reRNA_40b and reDNA) were purchased from biomers.net GmbH.

Table 1: Employed RNA and DNA sequenced

| Name | Sequence | Used in |
|---|---|---|
| miR-19b | 5' – UGUGCAAAUCCAUGCAAAACUGA – 3' | Fig. 2, 3, 4 and Suppl. Fig. 8, 11, 12, 14, 15, 18 |
| miR-19a | 5' – UGUGCAAAUCUAUGCAAAACUGA – 3' | Fig. 3 |
| miR-20a | 5' – UAAAGUGCUUAUAGUGCAGGUAG – 3' | Supp. Fig. 16 |
| miR-197 | 5' – UUCACCACCUUCUCCACCCAGC – 3' | Fig. 3 |
| reRNA_14b | 5' – 5AUGGC55AUGGC5 – 3' with 5 = 2'-OMe-A modification | Fig. 2, 3, 4 and Supp. Fig. 2, 6, 8, 11, 12, 14 - 19 |
| reRNA_40b | 5' – GGCUCAUUUGCAGGGGGGAGCCAAAAGGGUCAUCAUCUCC – 3' | Supp. Fig. 4, 6, 7 |
| reDNA | 5' – GCTGGGTGGAGAAGGTGGTGAA – 3' | Supp. Fig. 3, 5, 9, 10, 13 |
| RNU48 | 5' – GATGACCCCAGGTAACTCTGAGTGTGTCGCTGATGCCATCACCGCAGCGCTCTGACC– 3' | Fig. 3 |

For the preparation of the crRNA_19b, the construct was ordered as DNA from biomers.net GmbH with an additional T7 promoter sequence (marked in blue):

5' – AAGCTAATACGACTCACTATAGGGGGATTTAGACTACCCCAAAAACGAAGGGGACTAAAACTCAGTTTTGCATGGATTTGCACA – 3'

The crRNA DNA was annealed to a T7 primer and incubated with T7 polymerase overnight at 37 °C, using HiScribe T7 Quick High Yield RNA Synthesis Kit (New England Biolabs) as indicated by manufacturer. The crRNA was subsequently purified, using RNAClean XP beads (Beckman Coulter) as indicated by manufacturer. For the miR-20a, a crRNA matching the miR-20a sequence was designed accordingly.

**LwCas13a powered miRNA detection:**

Preliminary to the miRNA detection, the biosensor is pre-functionalized. The assay components (specified in table 2) are introduced to the chip, by applying 2 µl of the reagent to the inlet. Via capillary forces, the incubation area is filled until a hydrophobic stopping barrier is reached. After the incubation at 25 °C, the reagents are removed by applying a vacuum to the inlet, removing all unbound biomolecules. After a washing step, using 50 µl of wash buffer (10 mM PBS, 0.05% TWEEN® 20), the next assay component is applied.

**1) Off-chip miRNA targeting:**

For the off-chip targeting of miRNAs, all detection assays, unless otherwise stated, were performed in a standard tube of 500 µl containing the Cas13a reaction mixture: 125 nM purified Cas13a, 62,5 nM crRNA, 250 nM reRNA and 4 U $\mu l^{-1}$ murine RNase inhibitor (New England Biolabs) and varying amounts of target miRNAs in nuclease free assay buffer (40 mM Tris-HCl, 60 mM NaCl, 6 mM $MgCl_2$, pH 7.3). By placing the tube in an incubator at 37 °C, all reactions were allowed to proceed for 3 h, before introducing the mixture to the pre-functionalized biosensor chip. Following the incubation protocol (table 2), the assay is subsequently read out by applying a 40 mM glucose solution to the chip with a flow rate of 20 µl $min^{-1}$.

Table 2: Incubation protocol of all employed assay components for the off-chip miRNA targeting

| No. | Reagent | Concentration | Incubation time | Supplier |
|---|---|---|---|---|
| 1) | Streptavidin | 800 µg $ml^{-1}$ | 1 h | S4762 from Merck |
| 2) | Bovine serum albumin (BSA) in 10 mM PBS | 1% | 1 h | P3688 from Merck |
| 3) | Applying the CRISPR/Cas13a reaction mixture to the biosensor chip for 15 min at 25 °C | | | |
| 4) | Glucose oxidase labeled anti-fluorescein antibodies (Ab-GOx) | 0.83 µg $\mu l^{-1}$ | 15 min | Kit: ab102887 from abcam SAB4600050 from Merck |

**2) On-chip miRNA cleavage:**

For the on-chip miRNA detection, the biosensor is completely pre-functionalized, by following the incubation protocol (table 3). The applied CRISPR/Cas13a reaction mixture is exchanged every hour during the incubation process. Afterwards, the assay is read out by applying a 40 mM glucose solution to the chip with a flow rate of 20 µl $min^{-1}$.

Table 3: Incubation protocol of all employed assay components for the on-chip cleavage

| No. | Reagent | Concentration | Incubation time | Supplier |
|---|---|---|---|---|
| 1) | Anti-biotin antibodies | 400 µg $ml^{-1}$ | 1 h | B3640 from Merck |
| 2) | BSA in 10 mM PBS | 1% | 1 h | P3688 from Merck |
| 3) | Reporter RNA (reRNA) | 250 nM | 15 min | biomers.net GmbH |
| 4) | Biotin | 2 mg $ml^{-1}$ | 15 min | B4501 from Merck |
| 5) | Glucose oxidase labeled anti-fluorescein antibodies (Ab-GOx) | 0.83 µg $\mu l^{-1}$ | 15 min | Kit: ab102887 from abcam SAB4600050 from Merck |
| 6) | Applying the CRISPR/Cas13a reaction mixture to the biosensor chip for 3 h at 37 °C | | | |

**Fabrication of the biosensor chip:**

For the biosensor a polyimide substrate, Pyralux® AP8525R (DuPont) is used after 1 h of copper etching. By using lift-off technology, the platinum (Pt) electrodes are realized. The resist ma-N 1420 (Micro Resist Technology) is spin coated onto the substrate at 3,000 rpm for 30 s, exposed to UV (ultra violet) light on an exposure unit (Hellas, Bungard Elektronik) and developed. After a physical vapor deposition step, the resist is removed, realizing 200 nm thick Pt electrodes. To precisely define the electrode areas, to electrically isolate them and to realize small wells to implement later on a stopping barrier, a 5 µm thick SU-8 3005 (MicroChem Corp.) layer is applied by spin coating at 4,000 rpm for 30 s. After UV light exposure and developing, the wafer is hard baked in an oven for 1 h at 150 °C.

To remove SU-8 residues on the Pt electrodes, a low frequency oxygen plasma is for 3 min at 300 W at room temperature (Tetra-30-LF-PC, Diener) applied. Using galvanic deposition, an Ag/AgCl on-chip reference electrode is realized by galvanic deposition. First, all contact pads are covered with an UV sensitive tape (1020 R, Ultron Systems Inc.), then the galvanic silver deposition is performed in an Arguna S solution (Umicore Galvanotechnik), employing a current density of −7.45 mA $cm^2$ for 10 min with a silver counter electrode. By chlorinating the silver layer in a 0.1 M KCl solution at +3.2 mA $cm^{-2}$ for 5 min, using a Pt counter electrode, the reference electrode is finalized.

To realize 500 µm width microfluidic channels, different exposed 63.5 µm thick Pyralux® PC1025 (DuPont) DFR layers are stacked on the substrate. The DFR layers are first exposed for 50 s to UV light on an exposure unit and then developed in a 1% sodium carbonate ($Na_2CO_3$) solution at 42 °C in an ultrasonic bath. The DFR layers are subsequently immersed into a 1% HCl bath for 2 min. After developed layers are aligned on the substrate and fixed onto the substrate, using a laminator (HRL 350, Ozatec).

To implement a hydrophobic stopping barrier, separating the immobilization area from the electrochemical cell, small drops of 3% Teflon® (AF 1600, DuPont) are dispensed into small SU-8 wells. As a last step, the channels of the biosensor are sealed with another DFR layer, realizing the in- and outlet. To prevent bending of the wafer, two DFR layers are stacked on the backside of the substrate. The wafer is diced into the individual biosensors with a pair of scissors and is hard baked at 160 °C for 3 h. For further information, we refer to the video journal paper of Bruch et al.[2].

### Cost estimation of the functionalized biosensor

For the estimation of the costs for the fabrication and functionalization of the biosensor, the in house used container sizes and prices of every step are used in the calculations. As these prices a based on relatively small container sizes, the price will strongly vary, depending on the quantity of manufactured biosensor chips. For the calculations, a batch of 4 wafers, each containing 130 biosensors, are used.

Table 4: Estimation of the costs for the fabrication and functionalization of one biosensor

| No. | Step | Reagent/Material used | Costs per biosensor |
|---|---|---|---|
| 1) | Substrate preparation | Pyralux AP8525 | 0.045 € |
| 2) | Lift-off for platinum deposition | ma-N 1420 | 0.024 € |
| 3) | Platinum deposition | Pt PVD | 0.480 € |
| 4) | Electrode isolation | Su-8 3005 | 0.067 € |
| 5) | Reference electrode | Ag/AgCl | << 0.001 € |
| 6) | Dry film photoresist | Pyralux PC1025 | 0.017 € |
| 7) | Surface functionalizing | Streptavidin (off-chip targeting) | 0.099 € |
| | | Anti-biotin antibody (on-chip cleavage) | 0.186 € |
| 8) | Blocking of the surface | BSA | << 0.001 € |
| 9) | Reporter RNA | 6-FAM and biotin labeled reRNA | 0.004 € |
| 10) | Enzyme labelling | Ab-GOx | 0.015 € |
| | | **Sum of the functionalized “off-chip targeting” biosensor chip** | **0.751 €** |
| | | **Sum of the functionalized “on-chip cleavage” biosensor chip** | **0.838 €** |

### Assay component optimization for the off-chip miRNA targeting:

Each assay component was optimized in terms of incubation time and concentration. If not otherwise stated, following incubation protocol for each assay was used (table 5). All incubation steps were executed at room temperature (25 °C), except the cleavage process of the Cas13a, which was performed at 37 °C for 3 h, before applying the mixture to the biosensor for 15 min at room temperature. Between each incubation step 50 µl of wash buffer was used.

Table 5: Incubation protocol of employed assay components, if not otherwise stated

| No. | Assay component | Concentration | Incubation time |
|---|---|---|---|
| 1) | Streptavidin in 10 mM PBS | 400 µg $ml^{-1}$ | 1 h |
| 2) | BSA in 10 mM PBS | 1% | 1 h |
| 3) | Reporter RNA (reRNA_14b) in 40 mM TRIS | 250 nM | 15 min |
| | Murine RNase Inhibitor in 40 mM TRIS | 4 U $µl^{-1}$ | |
| | LwCas13a in 40 mM TRIS | 125 nM | |
| | crRNA in 40 mM TRIS | 62.5 nM | |
| | Target miRNA in 40 mM TRIS | 0 nM | |
| 4) | Glucose oxidase labeled anti-fluorescein antibodies (Ab-GOx) in 10 mM PBS | 0.83 µg $µl^{-1}$ | 15 min |

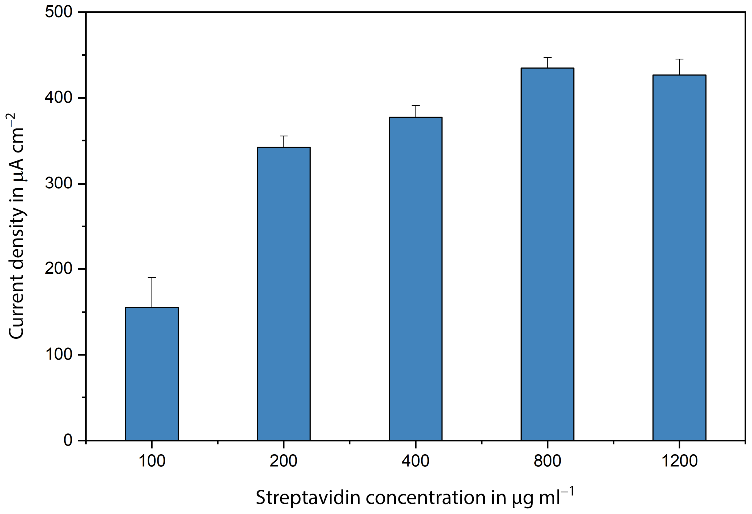

**Fig. 2 |** Comparison of different streptavidin concentrations to functionalize the surface of the chip. Error bars represent ± SD of n = 8 replicates.

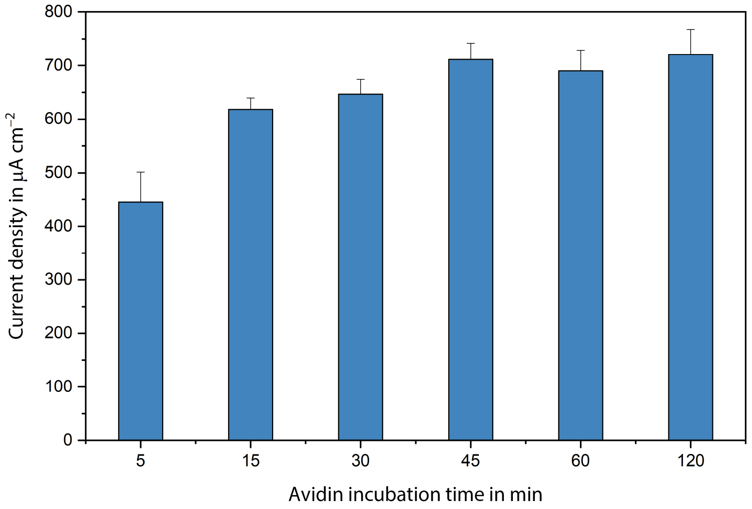

**Fig. 3 |** Investigation of the adsorption time of biomolecules, in this case avidin, to the channel surface. A concentration of 100 µg $ml^{-1}$ of avidin was incubated at 25 °C and blocked with BSA. A concentration of 1 µM biotin and 6-FAM labeled reDNA was applied and subsequently 16.67 µg $ml^{-1}$ of Ab-GOx was used for the coupling of the enzyme to the labeled reDNA. Error bars represent ± SD of n = 8 replicates.

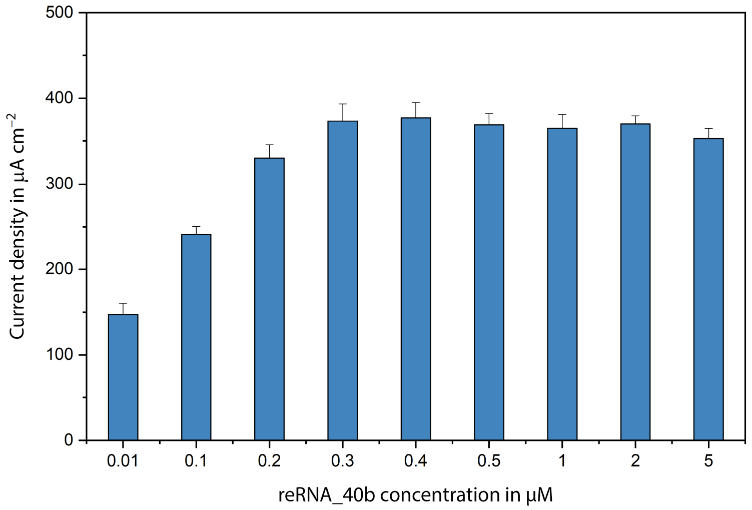

**Fig. 4 |** A surface coating of 400 µg $ml^{-1}$ of avidin was applied and blocked BSA. Different concentrations of the reporter reRNA_40b were applied and incubated in the channel for 15 min. Error bars represent ± SD of n = 8 replicates.

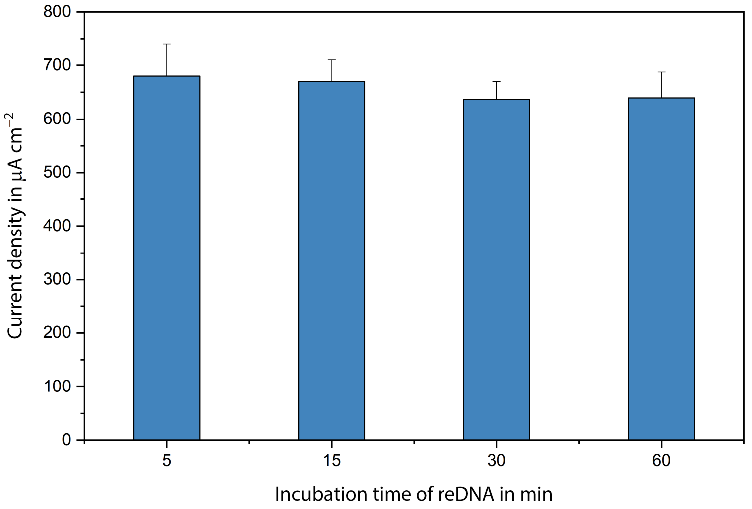

**Fig. 5 |** Investigation of the immobilization time of 1 µM of biotin and 6-FAM labeled reDNA after a surface coating of 100 µg $ml^{-1}$ of avidin and followed by 16.67 µg $ml^{-1}$ of Ab-GOx. Error bars represent ± SD of n = 8 replicates.

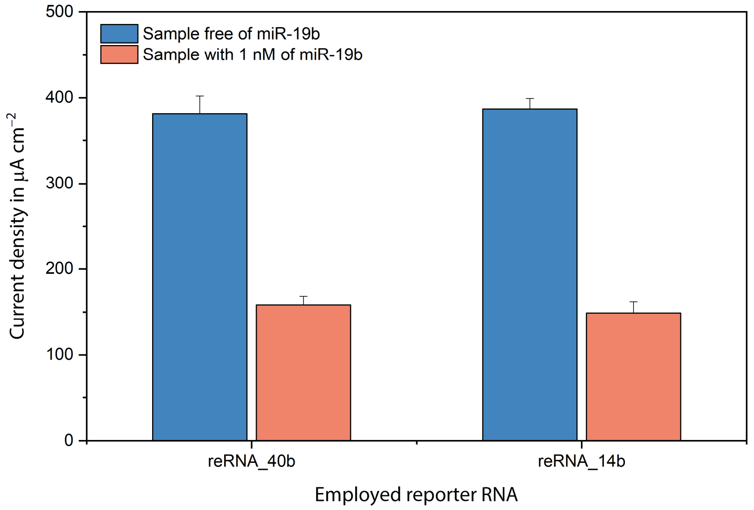

**Fig. 6 |** Comparison of two different reporter RNAs (reRNA_40b and reRNA_14b) after a surface functionalization with 400 µg $ml^{-1}$ avidin. The receptive reporter RNA is mixed with 144 nM of Cas13a, 72 nM of crRNA and 1 nM respectively 0 nM of miR-19b. Error bars represent ± SD of n = 8 replicates.

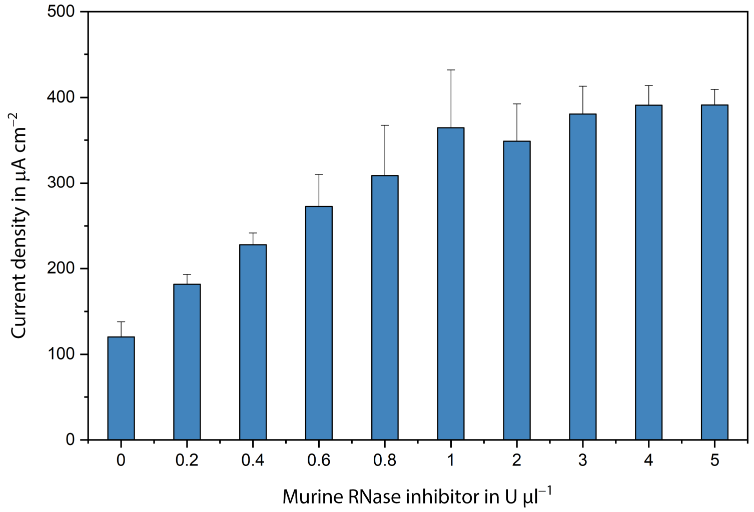

**Fig. 7 |** Optimization of the needed amount of murine RNase inhibitor. After an avidin (400 µg $ml^{-1}$) and blocking step (1% BSA) the inhibitor is mixed with the reporter RNA reRNA_40b (0.4 µM), the LwCas13a (144 nM) and the crRNA (144 nM) and incubated for 3h at 37 °C before introducing the mixture to the biosensor. Error bars represent ± SD of n = 8 replicates.

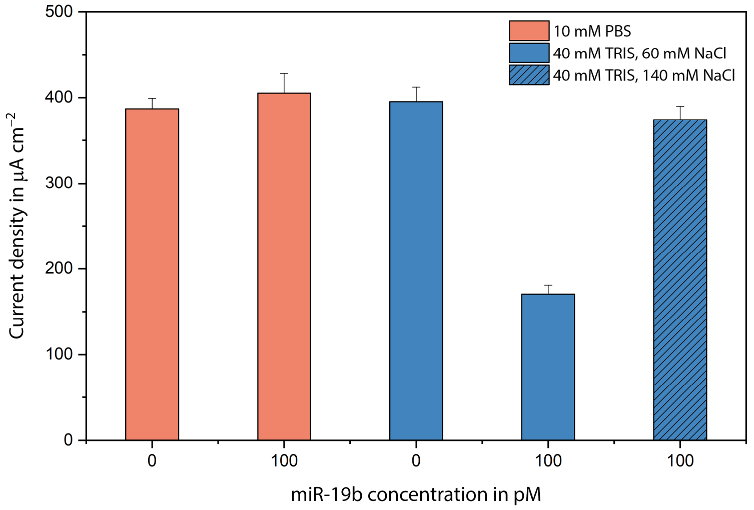

**Fig. 8 |** By mixing the LwCas13a, murine RNase inhibitor, crRNA and the sample (miR-19b) in different buffer (TRIS and PBS), the cleavage process is investigated. As a surface coating, 400 µg ml$^{-1}$ of avidin was used. From the left: error bars represent ± SD of n = 8 replicates for first three measurements and for the last two data points of n = 4 replicates.

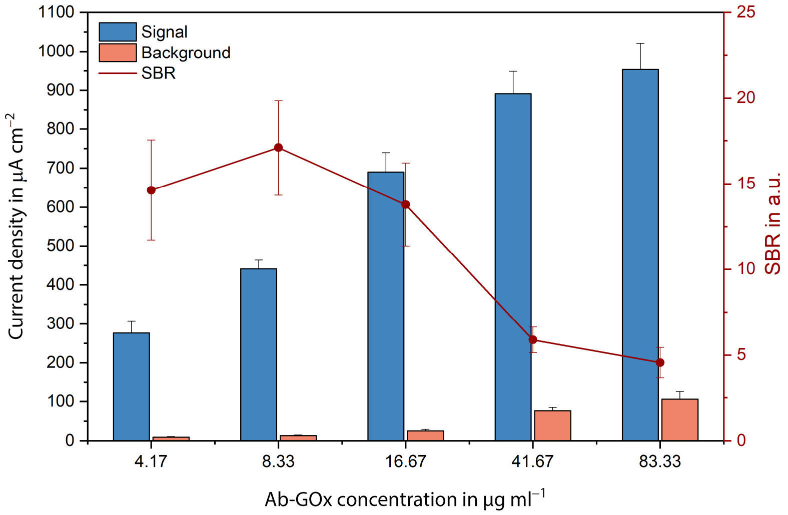

**Fig. 9 |** Dependency between the employed glucose oxidase labeled anti-fluorescein antibody (Ab-GOx) concentration, the background and the ratio of these two. After a surface coating of avidin (400 µg ml$^{-1}$), a blocking step (1% BSA) was followed. For the signal, 2 µM of biotin and 6-FAM labeled reDNA was incubated after which different Ab-GOx concentrations were applied. For the background signals, the double labeled reDNA is replaced by a biotin labeled reDNA with the same sequence. The signal to background ratio (SBR) was obtained by dividing the mean values of the signal by the mean values of the background. All error bars represent ± SD of n = 8 replicates.

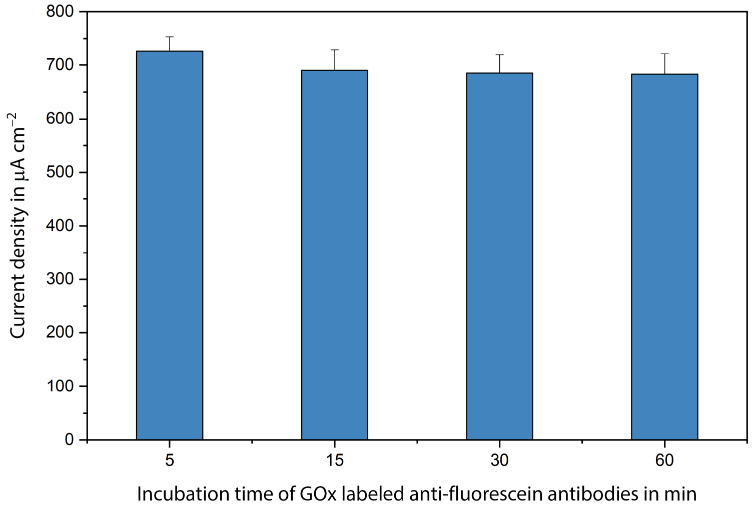

**Fig. 10 |** The incubation time of the GOx labeled anti-fluorescein labeled antibodies (16.67 µg $ml^{-1}$) were varied from 5 to 60 min, after an incubation of avidin (100 µg $ml^{-1}$) for 1 h, a BSA blocking step and the immobilization of 1 µM of biotin and 6-FAM labeled reDNA. All error bars represent ± SD of n = 8 replicates.

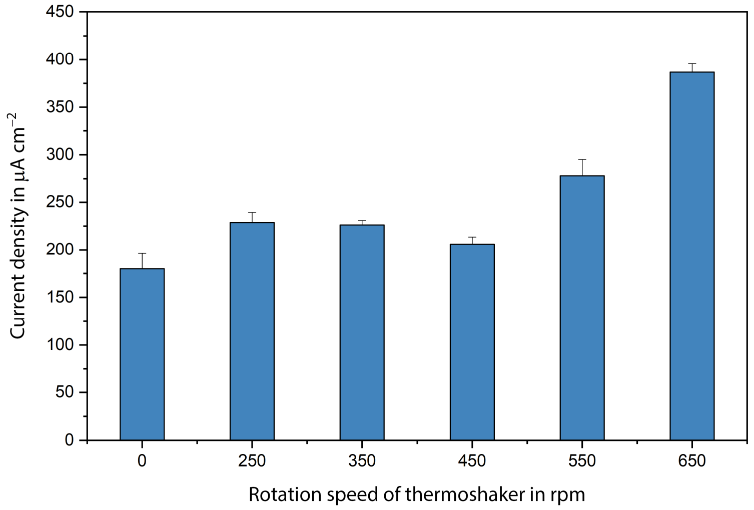

**Fig. 11 |** The dynamic cleavage process was performed on a thermoshaker (PST-60 HL plus from biosan) at 37 °C for 3 h with a miR-19b miRNA concentration of 50 pM. All error bars represent ± SD of n = 4 replicates.

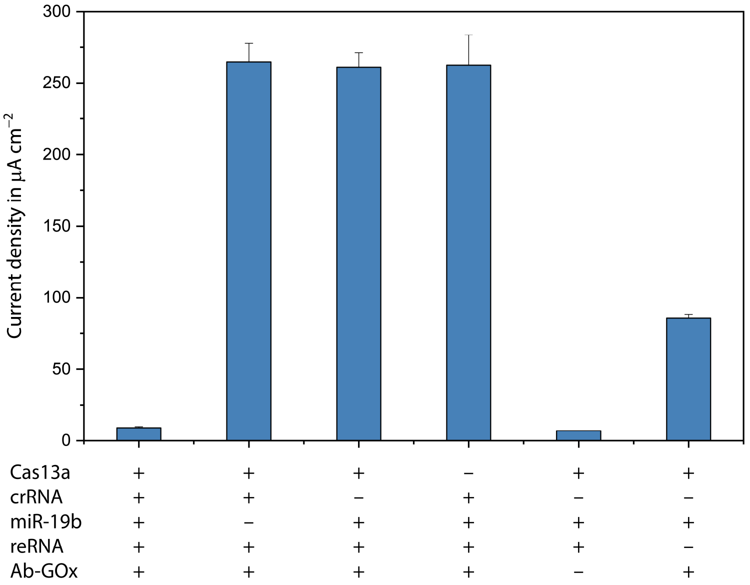

**Fig. 12 |** Negative Controls, using different reaction compositions, where '+' represents the presence and a '−' the absence of the reagent in the reaction volume. For the miR-19b, a concentration of 1 nM was chosen. All error bars represent ± SD of n = 4 replicates.

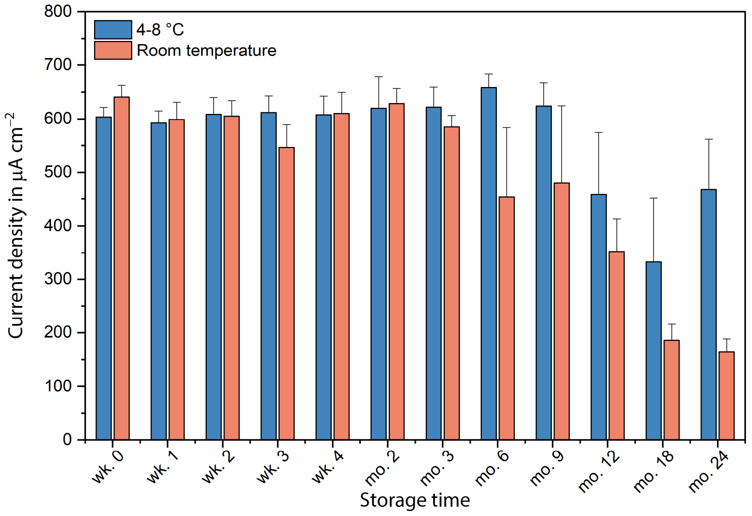

**Fig. 13 |** Long-term stability test by storing of pre-immobilized microfluidic biosensor chips (100 µg $ml^{-1}$ avidin for 1 h and blocking with 1% BSA for 1 h) at room temperature and at 4-8 °C in a standard fridge. After drying, by applying a vacuum to the inlet for 30 s, the chips are sealed in a bag, containing one silica pack. For the readout the chips were incubated with 1 µM reDNA for 15 min and with 16.67 µg $ml^{-1}$ of Ab-GOx. All error bars represent ± SD of n = 7 replicates.

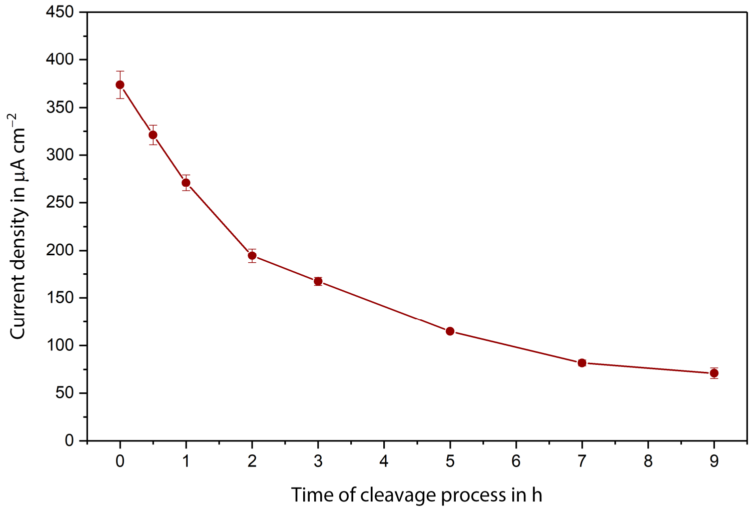

**Fig. 14 |** Time dependent off-chip cleavage process of a sample mixture, containing 50 pM of miR-19b. The solution was incubated for 9 hours with samples taken at several time points to perform the assay readout. All error bars represent ± SD of n = 4 replicates.

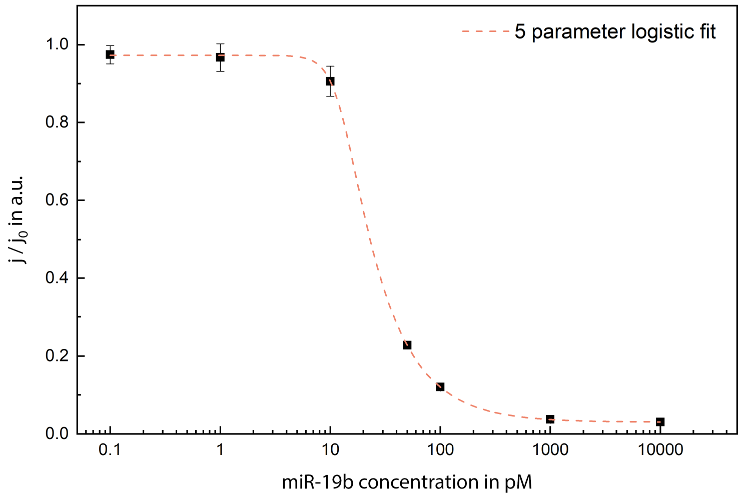

**Fig. 15 |** Calibration curve of the off-chip cleavage method, using the miR-19b as target miRNA. The incubation time for the cleavage process was 7 hours. The results are fitted with a 5-parametric logistic fit, resulting in a limit of detection of 10.5 pM. All error bars represent ± SD of n = 4 replicates.

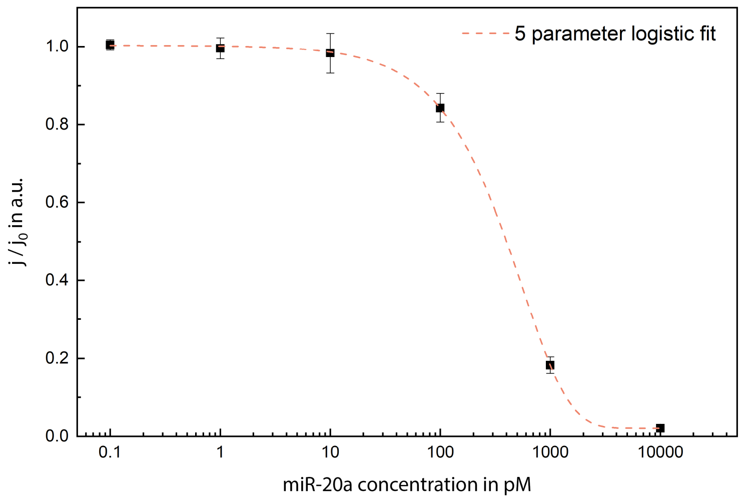

**Fig. 16 |** Calibration curve of the off-chip cleavage method, using the miR-20a as target miRNA. The incubation time for the cleavage process was 3 hours. The results are fitted with a 5-parametric logistic fit, resulting in a limit of detection of roughly 50 pM. All error bars represent ± SD of n = 4 replicates.

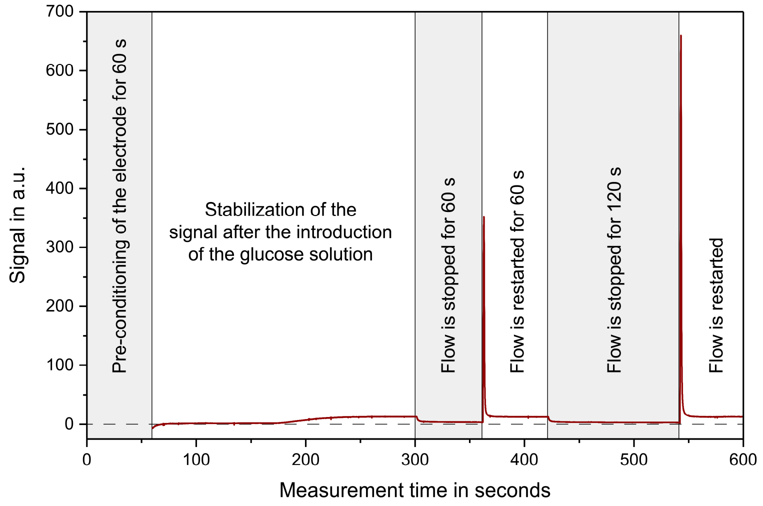

**Fig. 17 |** Readout procedure of our employed biosensor chip. After a 60 seconds step for the pre-conditioning of the on-chip working electrode, the glucose solution is introduced and the signal stabilizes over the next 4 minutes. The flow of the glucose solution is stopped for 1 minute, resulting in a first measurement peak and then restarted, followed by second stop for 2 minutes, resulting in the final measurement peak.

**Comparison of cleavage time**

In Table 6, all different calibrations curves are summarized. In general, the longer the incubation time for the cleavage process, the lower the limit of detection and the lower the dynamic range of the resulting calibration curve.

Table 6: Comparison of the different calibration curves in terms of LOD and dynamic range

| Cleavage time | Target | Limit of detection | Dynamic range |
|---|---|---|---|
| 1 h | miR-19b | 18 pM | 18 – 920 pM |
| 3 h | miR-20a | 50 pM | 50 – 3030 pM |
| 3 h | miR-19b | 10 pM | 10 – 820 pM |
| 7 h | miR-19b | 10 pM | 10 – 360 pM |
| 24 h | miR-19b | 2 pM | 2 – 50 pM |

**Serum sample preparation, RNA isolation and biochip measurement**

To isolate the RNA from the serum samples, the miRNeasy Kit from QIAGEN was used. The RNA was isolated from 100 µl of sample according to manufactures instructions. The eluate was mixed with the Cas13a, RNase inhibitor, crRNA and the reRNA and incubated at 37 °C for 24 hours and measured, accordingly to the in table 2 stated incubation protocol.

**Serum sample preparation, RNA isolation and real-time PCR**

The Isolate II Biofluids RNA Kit from Bioline was used for total RNA isolation from serum. RNA was isolated according to manufactures instructions and the final RNA concentration was measured with the nanodrop2000 system from Thermo Fisher Scientific. For the detection of the miRNA-19b and RNU48 the TaqMan system from Life Technologies was chosen. 10 ng of total RNA was applied for reverse transcription using the TaqMan MicroRNA RT Kit (Applied Biosystems, Life Technologies). The quantitative real-time PCR of miRNA-19b and RNU48 were performed in triplicates with the StepOnePlus System from Life Technologies. The miRNA levels were normalized to the stable internal control miRNA RNU48. Relative fold changes between expression of target genes in patient and control samples were calculated by using the $2^{-\Delta\Delta CP}$ method. The statistical analysis of the qPCR experiments was performed using the GraphPad prism 5 software.

**Assay component optimization for the on-chip miRNA cleavage:**

For the on-chip miRNA detection, the incubation protocol of table 3, excluding the use of biotin as an intermediate incubation step, is applied, if not otherwise stated.

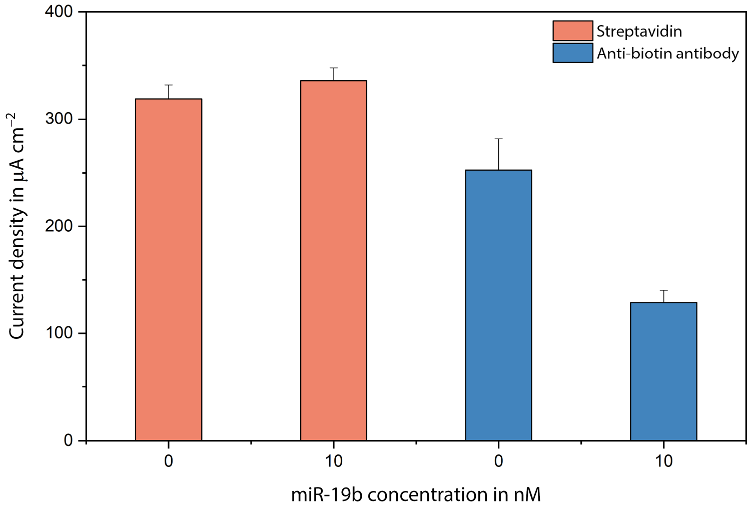

**Fig. 18 |** Comparison between a streptavidin (400 µg ml$^{-1}$) surface coating and the use of polyclonal anti-biotin antibodies (800 µg ml$^{-1}$) as a surface immobilization, each for 1 h of incubation. For both cases a concentration of 10 nM miRNA miR-19b was compared to a miR-19b free sample solution. All error bars represent ± SD of n = 4 replicates.

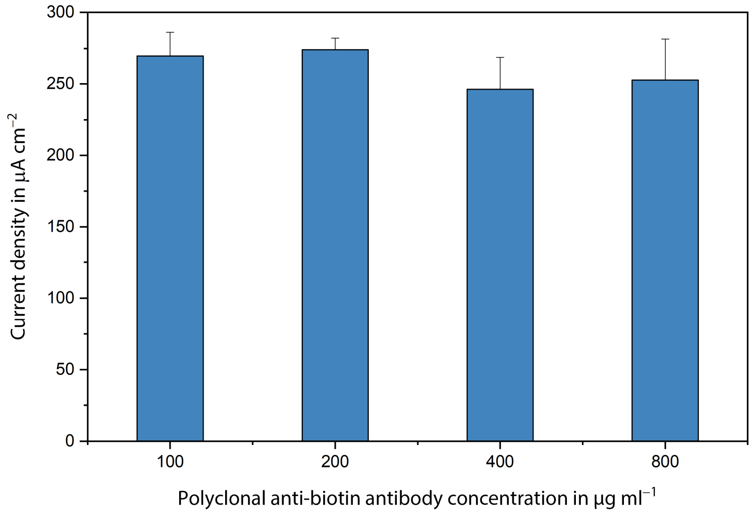

**Fig. 19 |** The concentration of the employed anti-biotin antibodies was varied from 100 to 800 µg ml$^{-1}$. All error bars represent ± SD of n = 4 replicates.

#### Future multiplexed approach of our biosensor

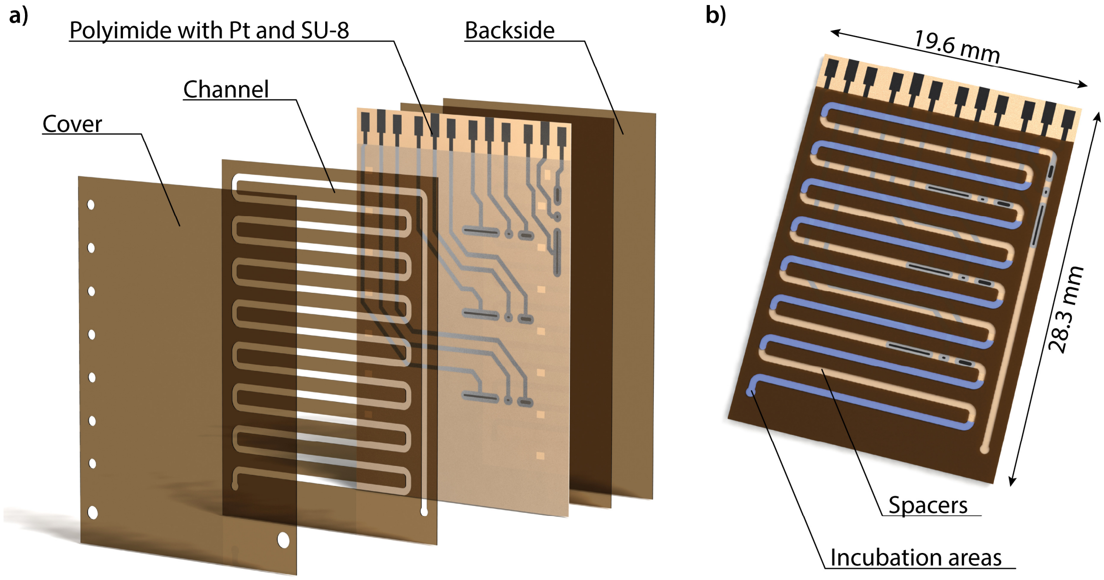

**Fig. 20 |** 3D Rendering of a possible future multiplexed biosensor for the detection of up to 8 miRNAs. **a)** Showing an exploded view of the biosensor with its different layers. **b)** Illustration of the functionality of the biosensor with its eight incubation (blue) and seven spacer areas. Each incubation area will be functionalized and can detect one specific miRNA and will be separated from other incubation areas by a biologically inactive spacer. One electrochemical cell will thereby detect two signals, which are time-wise separated.

#### Calculation of the limit of detection

For the calculation of the limit of detection (LOD) the fitted calibration curve is used to determine the signal response from the sensor at the LOD, using the following equation 1:

$$y = LOD_{Signal} = Signal_{blank} - 1.645 * SD_{blank} - 1.645 * SD_{lowest\_measured\_concentration}$$

To calculate the responding concentration of the obtained $LOD_{Signal}$, the equation of the 4 parameter logistic fitting curve is needed; equation 2:

$$y = A_{min} + \frac{A_{max} - A_{min}}{1 + \left(\frac{x}{x_0}\right)^p}$$

where $A_{min}$ is equal to the lower saturation plateau for higher concentrations, $A_{max}$ corresponds to the higher saturation plateau for very low concentrations. $x_0$ describes the concentration value for which 50% of the signal is present and $p$ is the slope of the fitting curve in the dynamic region. By placing the obtained $LOD_{Signal}$ of equation 1 into equation 2, the LOD can be calculated:

$$LOD_{concentration} = x_0 * \left(\frac{LOD_{Signal} - A_{max}}{A_{min} - LOD_{Signal}}\right)^{\frac{1}{p}}$$

These calculations were performed for every calibration curve, shown in this publication.

#### Overview of other electrochemical miRNA detection methods

Table 7: Other reported electrochemical miRNA detection methods based on cyclic voltammetry (CV), chronoamperometry (CA), differential pulse voltammetry (DPV) or square wave voltammetry (SWV)

| Detection method | Year | LOD | Target miRNAs | Employed techniques | Biological sample | Ref |
|---|---|---|---|---|---|---|
| In this work employed biosensor – CA | 2019 | 2.0 pM | miR-19b | CRISPR/Cas13a | RNA extracted from serum | - |
| Electrode surface blocking – DPV | 2018 | 1.0 pM | miR-21 | Magnetic beads for purification | RNA extracted from exosomes | 3 |
| Electrode surface blocking – SWV | 2013 | 5.0 aM | miR-21, 32, 122 | Three-mode sensor, protein based | Serum | 4 |
| DNAzyme based – SWV | 2016 | 5.2 pM | miR-155 | Reaction through displacement of strands | Spiked serum | 5 |
| Enzyme based – DPV | 2018 | 0.29 pM | miR-21 | Reaction through displacement of strands, rolling circle amplification | RNA extracted from cells | 6 |
| Enzyme based – CA | 2017 | 0.6 pM | miR-221 | Reaction through displacement of strands | RNA extracted from cells, spiked serum | 7 |
| Enzyme based – CA | 2016 | 0.14 fM | miR-155 | Graphene quantum dots | Spiked serum | 8 |
| Enzyme based – DPV | 2016 | 0.2 fM | miR-24 | Pd nanoparticles, opening of stem loop | - | 9 |
| Enzyme based – CV | 2014 | 1.0 fM | miR-141 | Tetrahedra DNA structure, opening of stem loop | - | 10 |
| Enzyme based – CA | 2014 | 40 pM | miR-21 | Magnetic beads | RNA extracted from cells | 11 |
| Enzyme based – CA | 2013 | 11 pM | let-7a | Opening of stem loop, secondary structure change | Diluted serum | 12 |
| Enzyme based – CA | 2013 | 7 pM | miR-222 | Magnetic beads | RNA extracted from cells | 13 |

**Notes:**

The University of Freiburg has filed for patent protection on the "Single-channel multianalyte biosensor" under the international publication number WO 2019/134741 A1. Dincer C., Kling A., Urban, G. A. and Bruch, R. are named as inventors on the patent[14].
